# Surface-Shaving Proteomics of *Mycobacterium marinum* Identifies Biofilm Subtype-Specific Changes Affecting Virulence, Tolerance and Persistence

**DOI:** 10.1101/2021.04.26.441561

**Authors:** Kirsi Savijoki, Henna Myllymäki, Hanna Luukinen, Lauri Paulamäki, Leena-Maija Vanha-aho, Aleksandra Svorjova, Ilkka Miettinen, Adyary Fallarero, Teemu O. Ihalainen, Jari Yli-Kauhaluoma, Tuula A. Nyman, Mataleena Parikka

**Author notes:** **Correspondence to:** Mataleena Parikka. These authors contributed equally to this work (Kirsi Savijoki, Henna Myllymäki). UoE Centre for Inflammation Research, Queen’s Medical Research Institute, University of Edinburgh, United Kingdom (Henna Myllymäki). Thermo Fischer Scientific, Ratastie 2, 01620 Vantaa, Finland (Adyary Fallarero).

## Abstract

The complex cell wall and biofilm matrix (ECM) act as key barriers to antibiotics in mycobacteria. Here, the ECM-proteins of *Mycobacterium marinum* ATCC927, a non-tuberculous mycobacterial model, was monitored over three months by label-free proteomics and compared with cell-surface proteins on planktonic cells to uncover pathways leading to virulence, tolerance, and persistence. We show that ATCC927 forms pellicle-type (PBFs) and submerged-type (SBFs) biofilms after two weeks and two days of growth, respectively, and that the increased CelA1 synthesis in this strain prevents biofilm formation and leads to reduced rifampicin tolerance. The proteomic data suggests that specific changes in mycolic acid synthesis (cord factor), Esx1-secretion, and cell-wall adhesins explain the appearance of PBFs as ribbon-like cords and SBFs as lichen-like structures. A subpopulation of cells resisting the 64 × MIC rifampicin (persisters) were detected in both biofilm subtypes, and already in one-week-old SBFs. The key forces boosting their development could include subtype-dependent changes in asymmetric cell division, cell wall biogenesis, tricarboxylic acid/glyoxylate cycle activities, and energy/redox/iron metabolisms. The effect of varying ambient oxygen tensions on each cell type and non-classical protein secretion are likely factors explaining majority of the subtype-specific changes. The proteomic findings also imply that Esx1-type protein secretion is more efficient in PL and PBF cells, while SBF may prefer both the Esx5- and non-classical pathways to control virulence and prolonged viability/persistence. In conclusion, this study reports a first proteomic insight into aging mycobacterial biofilm-ECMs and indicates biofilm subtype-dependent mechanisms conferring increased adaptive potential and virulence on non-tuberculous mycobacteria.

**IMPORTANCE:** Mycobacteria are naturally resilient and mycobacterial infections are notoriously difficult to treat with antibiotics, with biofilm formation being the main factor complicating the successful treatment of TB. The present study shows that non-tuberculous *Mycobacterium marinum* ATCC927 forms submerged- and pellicle-type biofilms with lichen- and ribbon-like structures, respectively, as well as persister cells under the same conditions. We show that both biofilm subtypes differ in terms of virulence-, tolerance- and persistence-conferring activities, highlighting the fact that both subtypes should be targeted to maximize the power of antimycobacterial treatment therapies.

## INTRODUCTION

Tuberculosis (TB) remains a major global health issue, with approximately 10 million new cases and 1.4 million deaths in 2019 (1). The causative agent, *Mycobacterium tuberculosis* (Mtb), is carried by an estimated one quarter of the human population as a latent infection, which has a 5–10% lifetime risk of developing into TB disease. In addition, the emergence of drug-resistant Mtb strains continues to be a public health threat, with about half a million new cases in 2019. Even in the case of drug-sensitive Mtb strains, the first line antibiotic treatment requires the use of four antimicrobials over a course of at least six months (WHO 2020). Moreover, despite successful treatment, the recurrence of TB carries a substantial risk, especially among immunocompromised patients (2, 3). The heterogeneity of the standard treatment outcome is also evident in PET-CT images showing non-resolving and active lesions and the presence of Mtb mRNA in sputum samples. This suggests that a significant proportion of patients generate viable mycobacteria in their lungs even after clinically curative antibiotic treatment (4). In a rabbit TB model, it was further shown that the caseum of granulomas contains Mtb that are highly tolerant to most anti-TB drugs (5). The complex mycobacterial cell wall, involving capsule and outer/inner membranes connected by a dense mycolyl-arabinogalactan-peptidoglycan with high lipid levels, is the main barrier that protects the bacterial cells against drugs (6). While the mechanisms leading to drug tolerance in TB have remained poorly understood, biofilm formation was recently indicated as one of the strategies to increase viability, tolerance and persistence (7-10).

Biofilm formation is defined as adherent growth within self-produced extracellular matrix/ECM consisting of proteins, polysaccharides, and DNA/RNA, and it is the strategy bacteria use to escape the effects of antibiotics and host defense systems (11-13). Mycobacteria use phenotypically distinct biofilm subtypes for growth, which genetically and physiologically differ from the planktonic-type growth. These include (i) floating/pellicle-type biofilms (PBFs) at the air-liquid interface having an ECM rich in free mycolic acids (MAs) and with a frequent cord/ribbon-like appearance, while (ii) submerged-type biofilms (SBFs) show adherent growth on a solid substratum (11, 14-16). The capsule layer plays a vital role in triggering biofilm growth in mycobacteria, as cells cultured in the presence of Tween-80 (non-ionic surfactant) has been shown to detach the capsule layer and prevent the biofilm formation (17). Thus, this labile layer forming the first molecular interaction with the host/environment is likely to involve key factors contributing to persistence/adaptation and search of anti-TB targets. Although several studies on mycobacteria have pinpointed cellular pathways and proteins that affect the capsule/cell wall and the biofilm formation (9, 14, 17-25), systematic investigation of the factors that directly interact with the surrounding environment is necessary to be able to maximize the power of antimycobacterial treatment therapies.

*Mycobacterium marinum* (Mmr) has proven to be an excellent alternative model pathogen for slow-growing Mtb, as it allows for the investigation of TB-like chronic and latent infections in its natural host, the zebrafish (26-29). Cultured mycobacterial biofilms have been used to understand resilient bacterial phenotypes emerging in mycobacterial infections. However, the distinct phenotypic profiles associated with PBFs and SBFs, including marker proteins discriminating the two biofilm subtypes have remained poorly understood. To shed light on the specific attributes linking these biologically different biofilm subtypes to their phenotypes, we first cultured Mmr strain ATCC927 to create in vitro biofilms. These biofilms were then imaged using widefield deconvolution microscopy (WDeM) to investigate temporal effects on the biofilm architectures. Label-free quantitative (LFQ) proteomics was next used to uncover the ECM-proteome dynamics in maturing Mmr biofilms and to identify the cell surface proteins (proteome) on Mmr grown on Tween-80, a detergent known to prevent cells from clumping and forming a biofilm (17). The key proteome findings were validated by gene overexpression studies to indicate cellulose-dependent biofilm formation as well as biofilm killing assays to confirm the formation of persister cells in both biofilm subtypes. To the best of our knowledge, this is the first study monitoring mycobacterial ECM-proteomes over three months’ time as well as protein and morphological phenotypic markers for distinguishing defined biofilm subtypes.

## RESULTS

### SBFs and PBFs show distinct morphological characteristics

The kinetics of development and maturation as well as the morphology of mycobacterial PBFs and SBFs has been reported to differ substantially (8). Here, we first show that that Mmr forms PBFs at the air-liquid interphase and SBFs attached onto the bottom of the culture well under the same physiological in vitro conditions after two weeks of growth **(Fig. 1SA)**. The SBF subtype develops earlier (visible already after two days of culture) than the PBF, which was not clearly distinguishable before two weeks of growth. Next, we investigated the three-dimensional morphology of Mmr biofilms in more detail by culturing Mmr cells, carrying the pTEC27 plasmid with the tdTomato fluorescent marker gene (29), for two and three weeks to produce PBFs and SBFs, and analyzing the biofilms by widefield deconvolution microscopy (WDeM). **Figure 1** shows that Mmr forms organized, three-dimensional structures with distinctive, subtype-specific morphological features. For the SBF, the structures displayed a lichen- or moss-like appearance, having tens of microns high feature structures rising from the biofilm base after two weeks **(Fig. 1B)**. In comparison, the morphology of the PBF subtype was very different by the first time point, showing flat, ribbon-like structures without any protruding structures **(Fig. 1C)**. Defined, extensive structures in all dimensions, although less dense compared to those detected at the two-week-time point, were found for both biofilm subtypes also after three weeks of growth.

**Figure 1.**
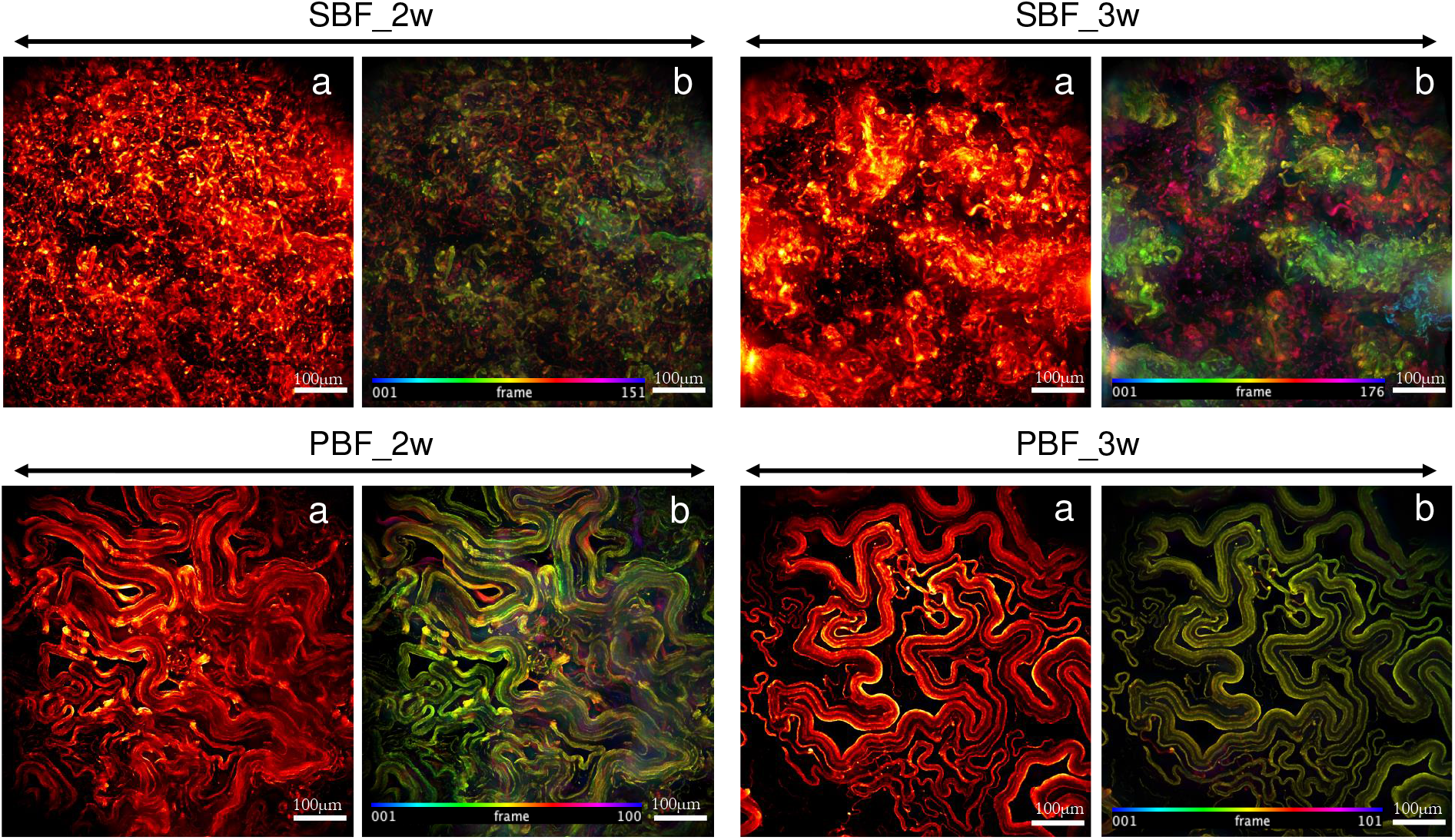
Mmr biofilms show distinct growth morphologies after two-weeks of growth. SBFs grow with lichen-like structures, whereas PBFs have a ribbon-like cords morphology, which becomes more defined with maturation (after 3 weeks). The WDeM images are maximum-intensity projections of one-, two-, and three-week-old biofilms (a) together with an image where Z-position is color-coded (b). X/Y scale bar corresponds to 10 µm and frame interval is 2µm.

### Submerged biofilms exhibit the greatest ECM-proteome diversity

As the phenotypic profiles of PBFs and SBFs are clearly different, their ECM-proteomes were next quantitatively monitored and compared during the development and maturation stages. To this end, the PBF and SBF cells at the points in **Figure 2A** were subjected to trypsin/Lys-C digestion as well as LC-MS/MS-based protein identification and LFQ (all data available via PRIDE with identifier PXD02010). Logarithmic state planktonic cells (PL_log), representing single-cell cultures, were obtained by growing the Mmr strain in the presence of Tween-80. The quality of each data set was high: 84.7% or all proteins were identified with at least three or more matching peptides, with an average sequence coverage of approximately 31% and only 11% of proteins were categorized as single-peptide-hits. In addition, a broad overlap in protein identifications was detected within the four biological replica samples; 41-89 % of the proteins were shared by each replicate, with the two-week PBF and the three-week SBF showing the highest variation between replicates **(Fig. S1B). Table S1** lists the proteins detected in at least two out of four replica samples. An outlier replicate associated with one of the SBF identification replica sets at the three-week-timepoint was excluded from subsequent data analyses. The number of detected proteins was 1132, 1957, and 2133 for the PL, PBF and SBF cells, respectively.

**Figure 2.**
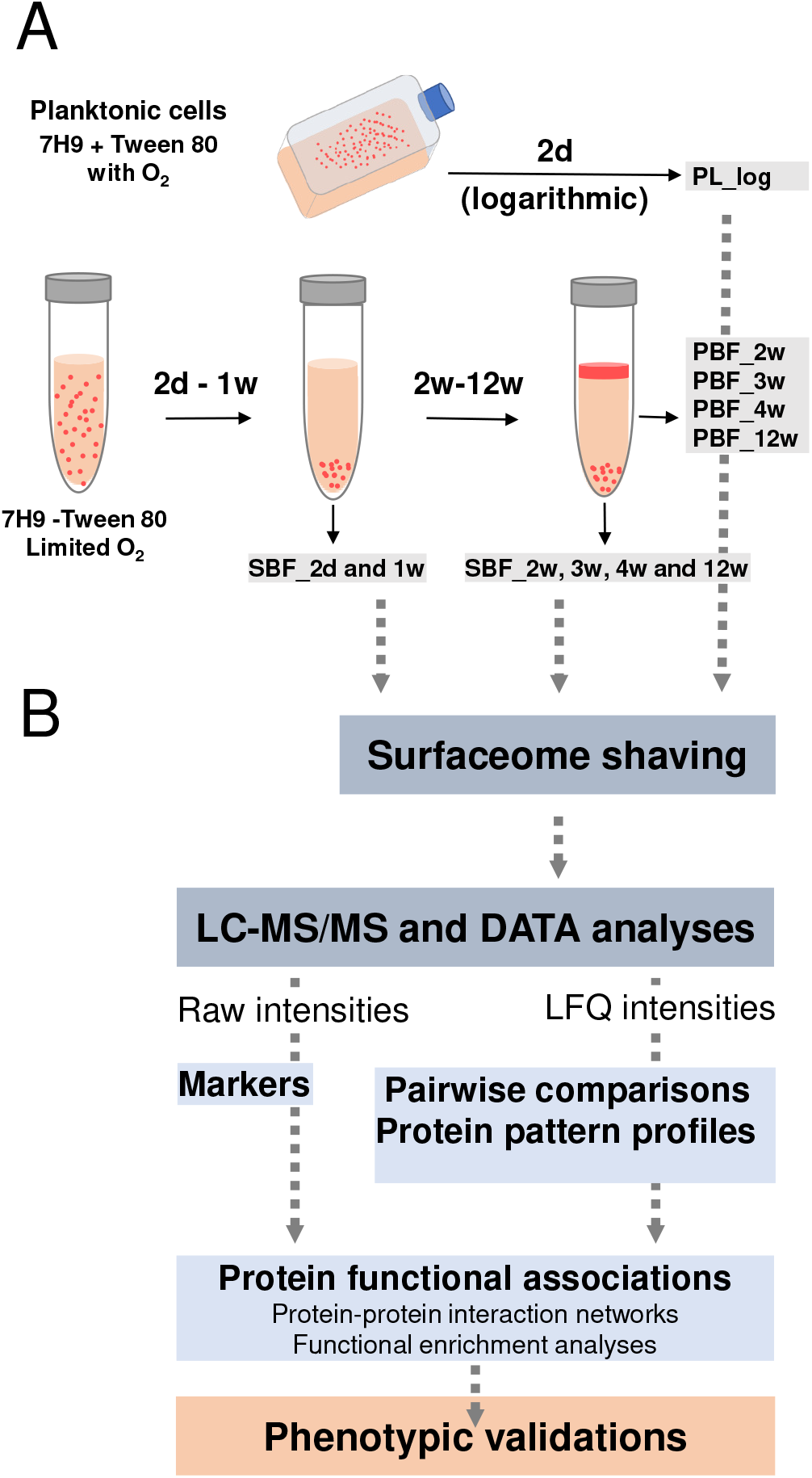
**(A)** A workflow depicting the conditions and time points used for preparing the planktonic and biofilm cells of Mmr. Grey arrows indicate sampling time points for pellicle (PBF) and submerged (SBF) biofilms. (B) Workflow used for identification of surface proteins associated with planktonic (PL_log) cells, PBFs and SBFs. Marker proteins were identified by comparing the raw intensity data, statistically significant protein abundance changes by pairwise comparisons of the log2 converted LFQ data and the protein co-abundance patterns by subjecting the LFQ data to imputation and z-score normalization. STRING and pathway enrichment analyses were conducted on the selected heat-map clusters and necessary phenotypic assays to validate the key proteome differences.

### Cytoplasmic protein export/release is most efficient in submerged biofilms

**Figure S2A** shows the distribution of all identified proteins according to their predicted secretion motif (Sec/SPII, TatP/SPI, LIPO/SPII, type VII secretion/T7SS, SecretomeP) and the number of transmembrane spanning domains (TMDs). The most notable differences were detected for membrane proteins with six to ten TMDs as well as in the number of cytoplasmic proteins. Nearly two-fold more TDM proteins were detected in the PL than the biofilm cells. In contrast, two-fold more cytoplasmic proteins predicted to be exported out of the cells via a non-classical route (SecretomeP) were identified from the biofilm-ECMs (*n*, 300) in comparison to the PL cells (*n*, 150). For many of these proteins, a secondary function as a moonlighting protein (30) could be indicated **(Table S1)**. In addition, over 900, 1600 and 1800 cytoplasmic proteins identified on the PL, PBF, and SBF cells, respectively, contained no motifs for classical or non-classical secretion and were assigned here as “Others” **(Table S1)**.

### Most significant protein abundance changes specific to planktonic and biofilm cells

The Venn diagram in **Figure S2B** indicates the highest number of specifically identified proteins on the SBFs (*n*, 173) and the lowest on the PBFs (*n*, 16), while no unique identifications were detected for the PL cells. The uniquely detected proteins with the highest raw intensity values included a signal transduction associated serine/threonine-protein kinase (PknL), a LGFP-repeat protein specific to SBFs, and a β-1,3-endoglucanase and bacterioferritin BfrA specific to PBFs **(Table S2)**. The proteins detected with the highest intensity values and only in the biofilm-ECMs included an error-prone polymerase DinB, a preprotein sec-translocase subunit YajC, a cytochrome P-450 monoxygenase, a PE family immunogen and a signal transduction-related adenylate cyclase involved in cyclic di-AMP biosynthesis **(Table S2)**.

Next, the log2 transformed MaxLFQ data was subjected to pairwise comparisons to indicate statistically significant protein abundance changes **(Table S3). Figure 3** shows the greatest growth mode- and time-dependent fold-changes related for the PL vs. biofilms cells, PBF vs. SBF cells and each biofilm subtype at different time points. Comparison of the PL and both biofilm cells at their first timepoints of growth (PBF_2w and SBF_2d) indicated the most prominent changes for PPE-family proteins (e.g., PPE61) and enzymes involved in cell envelope biogenesis/metabolism (MurE, CwlM, cutinase and CelA1). Among these, the PPE61 immunogen was *ca*. 6000- and 1800-times more abundant on the PL compared to the PBF_2w and SBF_2d, respectively. CelA1, a β-1,4-cellobiohydrolase known to prevent biofilm growth in *M. smegmatis* and Mtb (11, 18, 19), was detected with 50- and 130-fold higher abundances on the PL compared to the PBFs at the one-week and the SBFs at the two-day timepoints, respectively **(Table S3)**.

**Figure 3.**
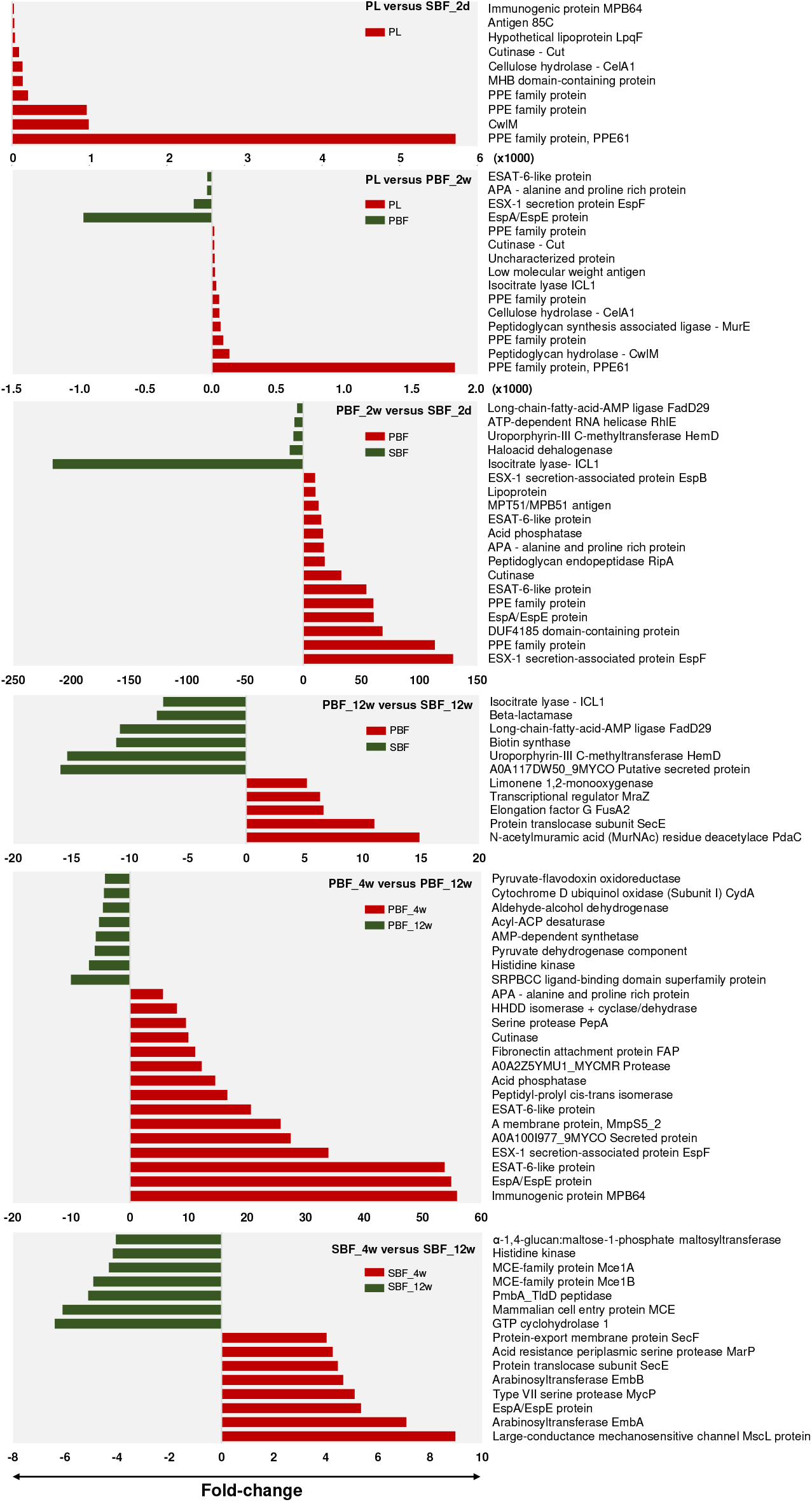
The most significant protein abundance fold changes between the indicated cell types at selected timepoints. The log2 transformed LFQ data was analyzed using Student’s t-test with permutation-based FDR adjustment. In the first two top panels, the fold-change x 1000.

Comparison of the PBF_2w and SBF_2d cells indicated Esx1-associated virulence factors (i.e., EspF, EspA/EspE and ESAT-6) and PPE family immunogens as 30–130-fold more abundant on the PBF than the SBF cells, meanwhile tricarboxylic acid (TCA)/glyoxylate cycle-associated isocitrate lyase (ICL1) was over 200-fold more produced by the SBF than the PBF cells. After 12 weeks, the proteins more abundant in the SBF compared to the PBF included an LppP/LprE lipoprotein (*ca*. 16-fold), HemD involved in the synthesis of vitamin B12 (*ca*.15-fold), FadD29 contributing to the synthesis of phenolic glycolipids (∼13-fold), β-lactamase able to hydrolase β-lactam antibiotics (*ca*. 9-times) and ICL1 catalyzing the glyoxylate shunt-mediated activities (*ca*. 8-fold). More abundant proteins on the PBF at this time-point were identified as a polysaccharide (N-acetylmuramic acid, MurNAc) deacetylase PdaC (*ca*.15-fold) and a translocase subunit, SecE (*ca*.11-fold).

In the PBF, an MPB64 immunogen, siderophore export accessory protein MmpS5, several Esx1-associated proteins (EspA/EspE, EspF) and adhesins (Ala-Pro-Ala rich protein APA and fibronectin binding protein FAP) displayed the most significant abundance decreases at the 12-week timepoint. In the SBFs, these proteins included a large-conductance mechanosensitive channel protein Msc, a membrane protein acting as the cells’ safety valve to relieve osmotic pressure, arabinosyltransferases EmbA/EmbB, the Esx1 associated EspA/EspE and the MycP1 protease. Proteins with the greatest abundance changes after 12 weeks in the SBFs included mammalian entry proteins (MCEs) and an α-1,4-glucan:maltose-1-phosphate maltosyltransferase.

### Decreased CelA1 synthesis is also required for biofilm formation in *M. marinum*

As our findings suggest that a lack of CelA could also promote the biofilm formation in Mmr, we tested this hypothesis by overexpressing the *celA1* gene in a Mmr strain equipped with pTEC27 with the tdTomato fluorescent marker (29). First, the *celA1* expression level in the transformed Mmr strain was confirmed by qPCR, indicating a *ca*. 150-times higher *celA1* transcription compared to the control strain carrying an empty pTECV27 **(Fig. 4A)**. Then we analyzed the morphology of both the SBFs and PBFs after two weeks using the CelA1-strain with WDeM. As seen in **Figure 4B**, the CelA1-strain showed altered morphology compared to the Mmr with pTEC27 (WT control strain). After two weeks of growth the CelA1-strain SBF showed a less defined/loss of the lichen-like morphology and lower total thickness when compared to the SBF control with pTEC27. Similarly, the CelA1 overproduction in Mmr resulted in disrupted and fuzzy ribbon-like cords associated with PBF-type biofilm growth, as the PBF cells with pTEC27 had well defined and tight ribbon like structures.

**Figure 4.**
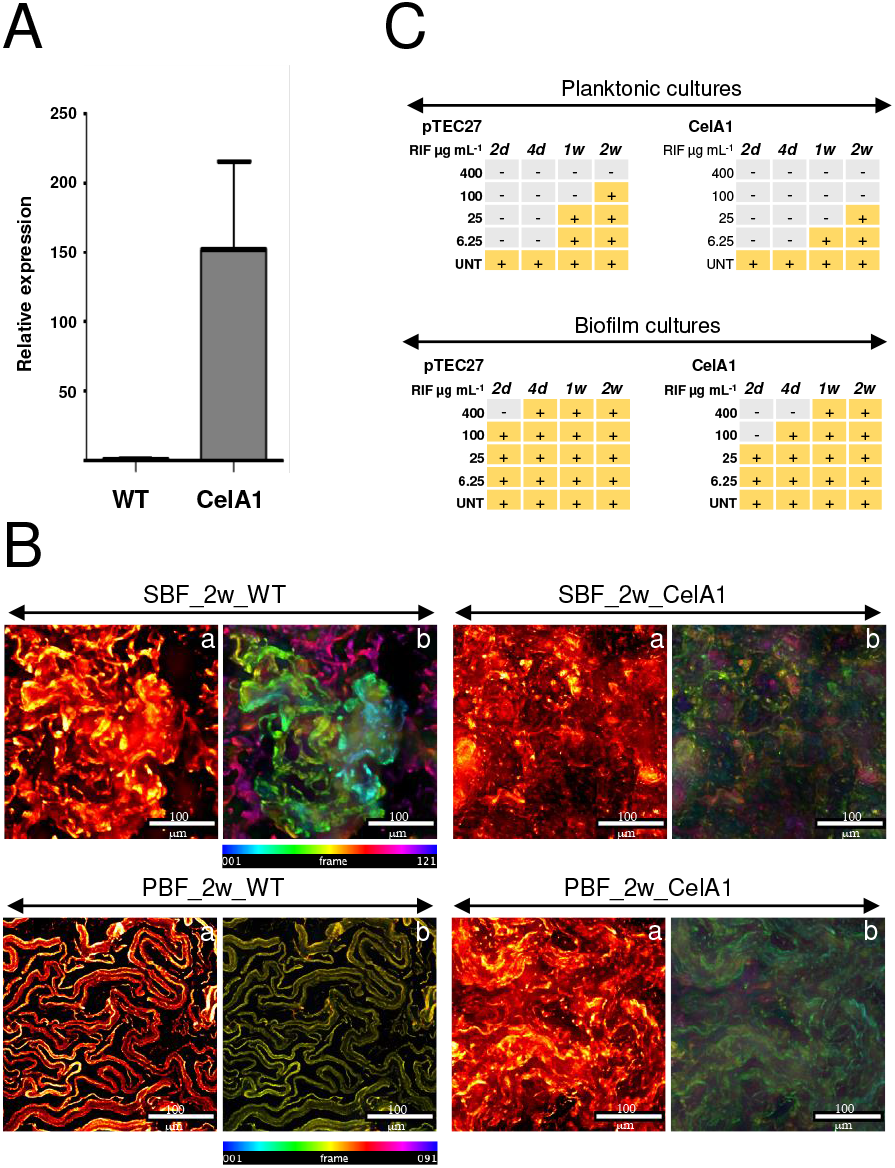
**(A)** Comparison of relative transcript abundance for *celA1* between the Mmr-CelA1 overexpression strain and the Mmr control strain with pTEC27 (WT). The CelA1 overexpression levels were normalized to the expression level of CelA1 in the WT control. The data is obtained from two technical replicates from two different bacterial clones. The bars represent the standard deviation. CelA1 expression is normalized with the amount of Mmar DNA in each sample. **(B)** CelA1 overexpression disrupts the biofilm development, and the formation of the subtype-specific growth morphologies. The WDeM images are maximum intensity projections of the two-week-old Mmr control biofilms (pTEC27, WT) and the Mmr-CelA1 cultures (a), together with color-coded by Z-position images (b). Scale bars 100 µm and frame interval is 2µm. **(C)** The MIC/MBC of rifampicin is reduced in both the PL and biofilms formed with the CelA1 overexpressing strain compared to the Mmr control cultures (PTEC27, WT). Rifampicin was added to the liquid cultures two, four, seven and 14 days after the start of the culture. Ten µl per sample (in triplicate) was plated seven days after the addition of rifampicin and CFUs were counted seven days thereafter. 100 CFU per sample was used as the cut-off limit for bacterial growth. The experiment was carried out three times. The figure shows a representative experiment. **-**, = no growth, + = bacterial growth, UNT= untreated

CelA1 expression was recently linked with biofilm formation, antibiotic tolerance, and virulence in Mtb (9). Therefore, Mmr cells in planktonic and biofilm forms with/without the CelA1 overexpression were also exposed to rifampicin to determine the minimum inhibitory (MIC) and minimum bactericidal concentration (MBC) for this bactericidal first-line TB drug. **Figure 4C** shows that, in both the planktonic and biofilm cultures, CelA1 overexpression decreases the MIC/MBC, with a clear impact on two-day-old and 4-day-old biofilms. These results indicate that CelA1 impedes biofilm formation and increases the susceptibility of the residing cells to rifampicin in Mmr.

### Functional pathways specifically induced in planktonic and biofilm cells

The LFQ proteomic data was next subjected to a PCA analysis for comparing growth mode- and time-dependent protein abundance patterns on the PL cells and aging biofilms. The PCA in **Figure 5A** shows clear clustering for each data set except for replicates associated with two-week-old PBF-proteomes, which show greater variation. PC1, separating the samples according to the growth mode, explains 39% of the total variation, while 24% (PC2) of the variation can be explained by the age of the culture. The two-day-old SBF-proteomes form a clearly distinguishable cluster, while the PL-proteomes and proteomes associated with the PBFs between the two- and four-week time-points show close clustering. Although the SBF- and PBF-proteomes differ greatly within the first four weeks of growth, these biofilm subtypes seem to undergo similar proteome changes during the later stages of growth, as proteomes of both subtypes clustered more closely at the 12-week timepoint. Notably, PBFs during the first weeks (two to three weeks) of growth shared a more similar ECM-proteome with the PL cells than the SBFs under the same conditions.

**Figure 5.**
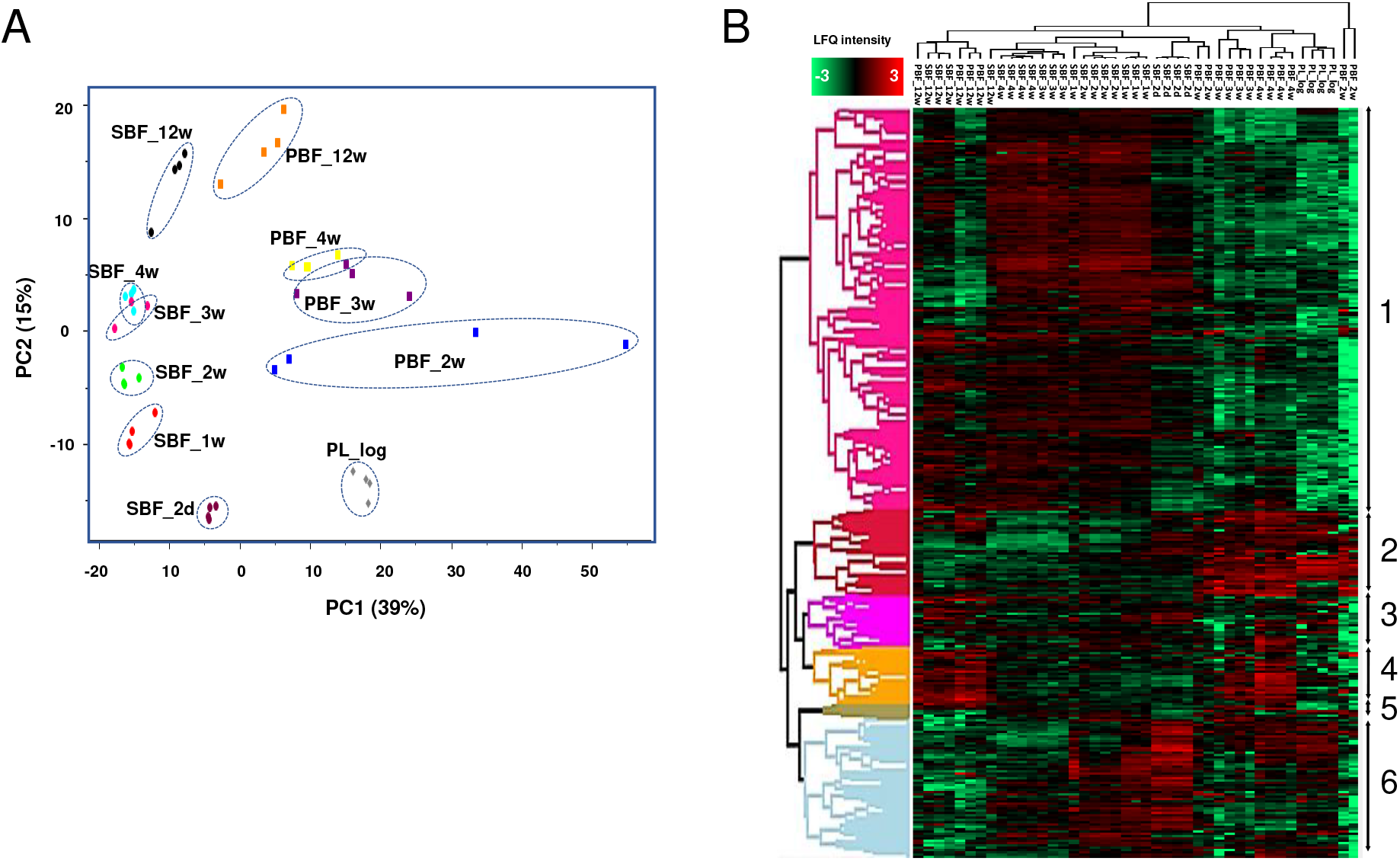
**(A)** A PCA analysis of all detected proteins (based on LFQ intensities excluding one SBF_3w outlier) with PC1 and PC2 indicating growth mode- and time point-dependent changes. **(B)** Hierarchical clustering of proteins (complete linkage; n, 690) with significantly changed expression profiles. Color intensity: red and green indicate higher and lower protein abundances, respectively.

Next, a multi-sample test (ANOVA) was conducted on the normalized LFQ intensity data to investigate growth mode-dependent proteome differences at time points between two days and three months. A dendrogram/ heatmap in **Figure 5B** shows hierarchically clustered co-abundance data for 690 proteins having a statistically significant abundance change in at least one of the conditions tested **(Table S4)**. Six major clusters were clearly distinguished, among which cluster 1 (*n*, 375) and cluster 6 (*n*, 125) contained the greatest number of proteins, with higher abundances in one-to four-week-old SBFs (cluster 1) and two-day-to two-week-old SBFs (cluster 6), respectively. STRING (Search Tool for the Retrieval of Interacting Genes/Proteins) enrichment analyses performed on both clusters **(Table S5)** indicated the greatest changes for pathways coordinating cell envelope biogenesis/ metabolism, energy metabolism and protein secretion/export. **Figure 6A** shows a protein-protein interaction (PPI) network for cluster 1 proteins: **(i)** cytoplasmic proteins with a primary function in amino acid biosynthesis (*e*.*g*., Gly, Asp, Tyr, Arg, His, Thr, Ser, Lys, Phe), purine/pyrimidine metabolism (*e*.*g*., PyrG, PurD/L/H, GuaB) and stress response (HrcA, ClpC/X, DnaJ, HtpG, AhpC, SodC, RecA, Trx), **(ii)** proteins involved in cell-wall/outer layer and mycomembrane biogenesis/metabolism (*e*.*g*., PknA/B, Weg31, CwsA, CwlM, PbpA1a, EmbA/B, KasA, DesA1/2, PpsA/B/D, PcaA, Fad enzymes), **(iii)** components of the respiratory electron transport chain (SDH, FMR, Qcr-complex) and ATP synthesis (F1F0 ATP synthase-complex), and **(vi)** proteins involved in iron storage/homeostasis (ferritin). The PPI network analysis on the cluster 6 proteins indicated the enrichment of metabolic activities related to translation (ribosomal proteins/r-proteins), stress response (GroEL/ES, GrpE, DnaK, TF, ClpB) and the TCA/glyoxylate cycle (*e*.*g*., CitA, ICL1, FBA, GlcB) **(Fig. 6B)**.

**Figure 6.**
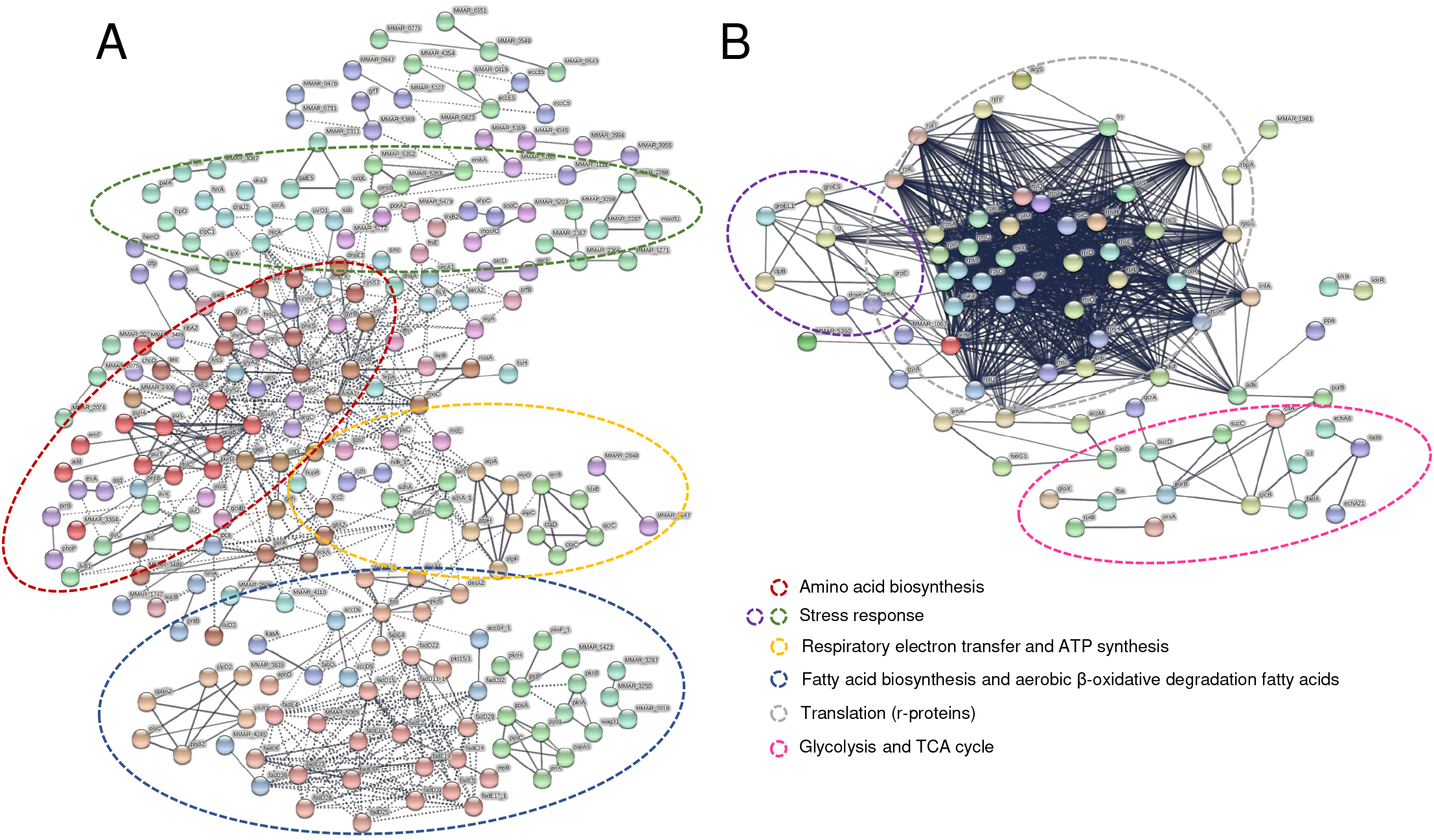
**(A)** A PPI network analysis on cluster 1 proteins (Fig. 5B) with higher abundancies on SBFs between two and four weeks. No. nodes, 368; no. edges, 3256; PPI enrichment *P* < 1.0e-16. **(B)** A PPI network analysis on cluster six proteins with higher abundancies on SBFs between two days and 2 weeks. Proteins were clustered using MCL with the inflation parameter set to 4.0 (cluster 6) and 6.0 (cluster 1). No. nodes, 155; no. edges, 3024; PPI enrichment *P* < 1.0e-16. Circles indicate the most enriched protein interactions.

Clusters 2, 4 and 5 (*n*, 144) share co-abundance patterns, which indicate increased protein abundances during the first weeks of growth in the PBFs when compared to the SBFs. These contain virulence-, invasion- and viability/persistence-related proteins, such as EsxA/B, ESX-EspB/G/M/P/N, Esx5-secretion associated protease MycP, cutinase (Cut), a lysophospholipase (YtpA), endopeptidase (Lon), heparin binding hemagglutinin (HbhA), fibronectin binding (Apa), catalase-peroxidase (KatG), and mammalian entry proteins (MCEs). Cytoplasmic proteins were also detected in these clusters (e.g., ICL2, ACN, ENO, GapDH, GPD, Tpi, PGK, MDH, ClpP1/2, CpsA/D, Trp, Cys, Met, an 18-kDa β-CA) but their composition differs clearly from those in clusters 1 and 6. In addition, cluster 2 contains virulence-associated ESAT-6-like proteins, TDM-cord factor synthesis associated Ag85A/C (mycolyltransferases), and an MPT64 immunogen with higher overall abundancies on the PL and PBF cells compared to the SBFs. The remaining cluster 3 (*n*, 47) differs from the other five by proteins with its overall higher abundancies on the PL cells and/or on four-to 12-week-old PBFs when compared to the SBFs at the same timepoints. One of these was identified as a potential trehalase (A0A2Z5YJK7_MYCMR), a glycoside hydrolase that catalyzes the conversion of trehalose to glucose, which had a high abundancy in four- and 12-week-old PBFs.

The protein identifications most relevant to biofilm growth and viability identifications are listed in **Table S6** according to their predicted functions. The major growth mode-dependent changes associate with the following five functional groups: **(i)** secretion mechanisms, virulence, and adherence; **(ii)** cell wall/membrane/lipid biogenesis and metabolism and biofilm formation; **(iii)** stress response; **(iv)** TCA/glyoxylate cycles and carbohydrate metabolism; and **(v)** maintaining redox balance and energy metabolism. An additional schematic model of the mycobacterial cell envelope in **Figure 7** illustrates the key proteome changes relevant to the PL-, SBF- and PBF-type growth in Mmr.

**Figure 7.**
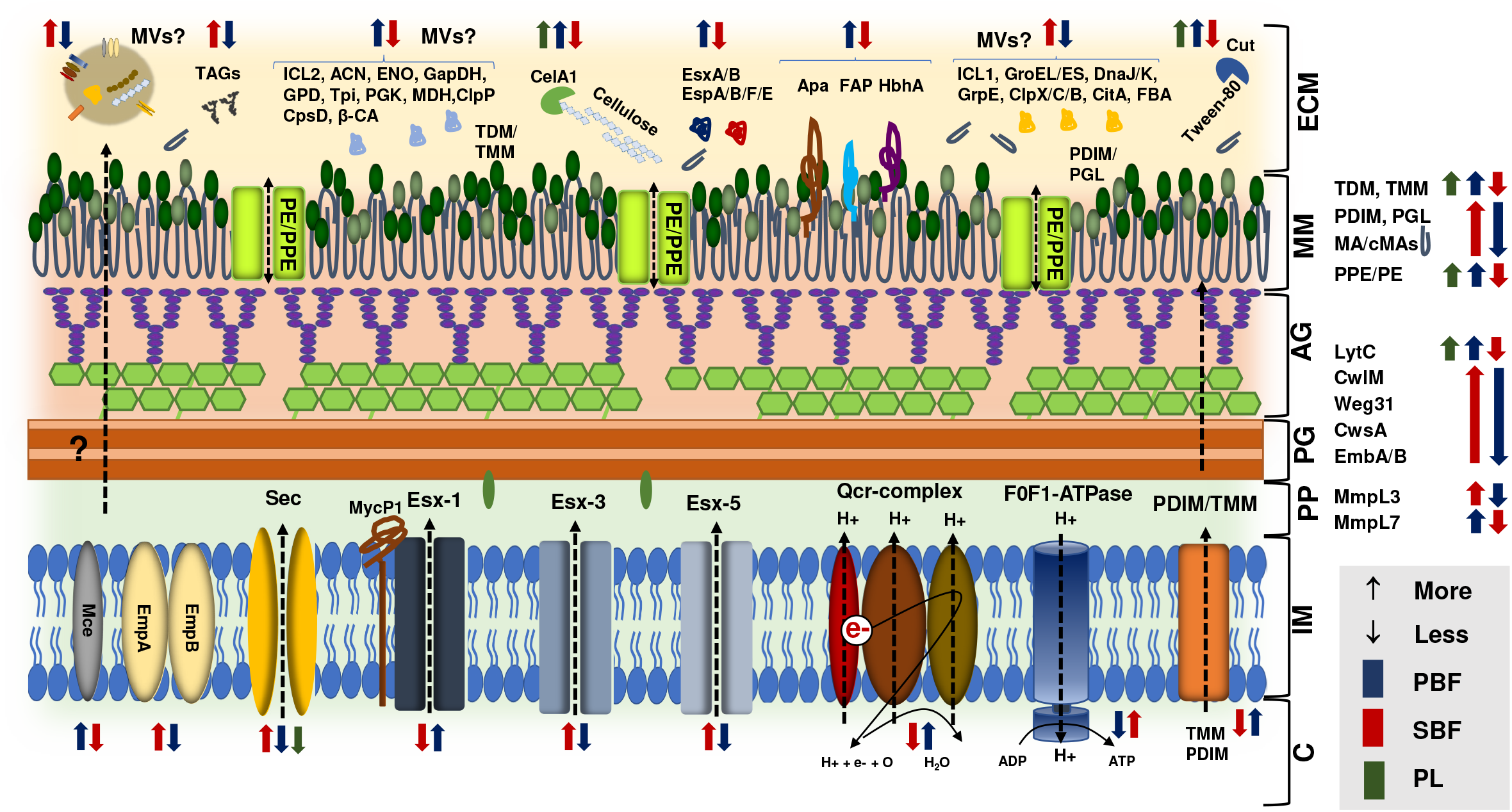
A schematic model of the Mmr cell envelope with key protein abundance changes specific to PL, PBF and SBF cells. Colored arrows pointing up/down refer to protein abundances/abundance changes within the indicated cell sample types (green, PL; blue, PBFs; red, SBFs). MA, mycolic acids; cMAs, cyclopropanated mycolic acids; TDM/TMM, trehalose-6,6-dimycolate/trehalose monomycolate; PDIM/PGL, phthiocerol dimycoceros-ates/phenolic glycolipids. C, cytoplasm; IM, inner membrane; PP, periplasmic space; AG, arabinogalactan; PG, peptidoglycan; MM, mycomembrane; ECM, extracellular matrix.

### Time-kill curve analysis for indicating persister cells in maturing biofilms

As growth mode-dependent differences imply higher persistence/tolerance-associated activities in biofilms than in planktonic cultures, we next validated these findings by exposing both the planktonic and biofilm cells to rifampicin and monitored cell death using a time-kill curve analysis. This method enables the demonstration of an overall slower killing efficacy for tolerant populations or a bimodal time-kill curve that indicates the presence of a persistent bacterial subpopulation (31, 32).

First, we used a bacterial killing assay with bioluminescence as a readout to quantify the tolerance/persistence in the planktonic cultures and two-week-old biofilms. The planktonic and biofilm cells were treated with 400 μg mL^-1^ rifampicin (64 x MIC, minimum inhibitory concentration), and the rate of bacterial killing was monitored for seven days. The use of bioluminescence as a readout for killing biofilm-associated bacteria was also assessed using an OD_600_-based method **(Fig. S3A)**. The time-kill curve for the biofilm population was bimodal, showing the faster killing of a susceptible subpopulation followed by a slower killing of a persistent subpopulation of cells (**Fig. 8A**). These results indicate that Mmr biofilms harbor significantly more persister cells than logarithmic phase planktonic populations.

**Figure 8.**
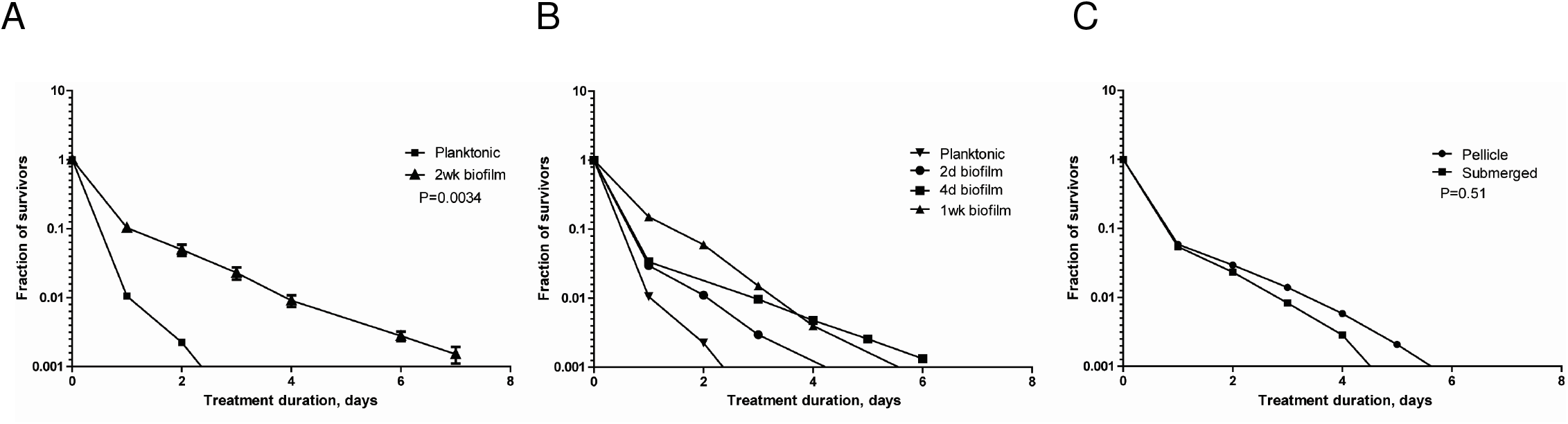
The proportion of persistent bacterial cells increases in Mmr biofilms. Time-kill curve analysis was performed by culturing biofilms from two days to two weeks and treating the bacteria with 400 μg mL^-1^ rifampicin. The killing kinetics were monitored for seven days by measuring the bioluminescence signal produced by Lux-Mmr daily. **(A)** Logarithmic growth phase planktonic and 2-week-old biofilm Mmr were treated with 400 µg mL^-1^ of rifampicin. The time-kill curves of the planktonic and biofilm-associated bacteria were significantly different (*P* < 0.0034, log-rank test). The means and SEMs of three biological replicates are shown. **(B)** In biofilms, persistence increases over time and is significantly higher after one week compared to planktonic bacteria (*P* = 0.0002, log-rank test). Planktonic culture and two-day-old biofilm show similar killing curves. Means and SEMs of three biological replicates are shown. **(C)** Two-week-old PBFs and SBFs were tested separately for persistence. The two different biofilm-types show no difference in their persistence levels (*P* = 0.51, log-rank test). Means and SEMs of three biological replicates are shown.

Next, the development of persistence in the biofilms was monitored by killing two-day-, four-day-, and one-week-old biofilm cells with 64 x MIC rifampicin. Analysis of the time-kill curves showed that persistence increases gradually in the maturing biofilms, reaching a statistically significant increase in one-week-old biofilms compared to the planktonic cells (*P* = 0.0002) (**Fig. 8B)**. In untreated biofilms, the bioluminescence signal level continues to increase well past the one-week timepoint, showing that the biofilm-associated mycobacterial population is replicating and/or metabolically active at this stage **(Fig. S3B)**. This indicates that increased persistence is not (mainly) caused by the induction of dormancy or metabolic inactivity. According to our experimental settings, PBFs form later than SBFs and are visually detectable only after two weeks. Thus, this data shows that a substantial persister subpopulation develops in SBFs by the first week of biofilm development.

To test if the formation of persister cells differs between the two biofilm subtypes, PBFs and SBFs were collected separately and tested with the time-kill assay under 64 x MIC rifampicin. After seven days, the time-kill curves indicated no significant differences in the rate of persistence between the two-week-old pellicle and submerged biofilms (*P* = 0.51) (**Fig. 8C**). Thus, our results indicate that the proportion of persisters is greater in over one-week-old Mmr biofilms than in logarithmic planktonic cell populations, and that the biofilm-associated persistence increases over time.

## DISCUSSION

### Mmr grows in morphologically distinct biofilm subtypes *in vitro*

A recent study confirmed that Mtb forms biofilm-like communities in vivo, which confers increased tolerance to rifampicin and thus provides an explanation for the chronic nature of TB (11). The present study shows that Mmr grows in two different biofilm subtypes, and that reduced CelA1 hydrolase activity is one of the main triggers of biofilm growth and increased tolerance to rifampicin in both biofilm subtypes. Studies on Mtb and *M. smegmatis* have demonstrated that cellulose filaments are vital structural constituents of mycobacterial biofilm-ECMs as well as essential for biofilm formation and the development of tolerance/persistence (9, 11, 18, 19). We also show that the Mmr biofilm subtypes show distinct morphologies, with SBFs containing lichen-like structures and PBFs consisting of ribbon-like cords under the same in vitro conditions. Biofilm growth accompanied by cording-like growth morphology is also reported for other mycobacteria and Mtb, in which the surface interactions mediated by e.g., mycolic acids modulating the mycomembrane/capsule hydrophobicity (11, 33). The proteomic data presented here suggest that subtype-specific changes in cord-factor TDM-synthesis (mycolyltransferase Ag85), Esx1-secretion, phthiocerol dimycocerosate (PDIM) export (MmpL7), MA cyclopropanation (PcaA/Cma2), and lectin synthesis (33-37) may have affected the mycomembrane composition and thereby contributed to distinct biofilm growth morphologies in Mmr.

### Mmr may use membrane vesicles to deliver proteins in the biofilm-ECM

The LFQ proteomics identified cytoplasmic proteins and proteins associated with the inner-/mycomembrane as the largest protein group in both the planktonic and biofilm cells. These findings are supported by studies identifying cytoplasmic proteins in the capsule of another Mmr strain (E11) and by showing that their number increases when mycobacterial cells grow in the biofilms, as demonstrated for *M. bovis* (17, 20). Membrane vesiculation is the most likely explanation for their presence on Mmr cells and within the biofilm-ECM, as several reports have demonstrated the presence of MVs on mycobacterial cells (38) as well as trapped in biofilm-ECMs in other bacteria (39). In addition, several of the cytoplasmic and inner-/mycomembrane-proteins detected here, including e.g., enzymes involved in cell wall synthesis and lipid/ fatty acid metabolism, were previously identified in MVs released by *Mycobacterium avium* 104 in response to starvation (40). Mycobacteria have been shown to form MVs from mycomembrane (mMV) during normal growth (cell lysis/death) and/or from inner-membrane (iMV) by blebbing in response to stress (e.g., iron-limitation and anoxia) (38). This report supports the idea that the identified myco-/inner-membrane-proteins could have also entered the biofilm-ECMs by MVs in our study. We further propose that CwlM, a N-acetylmuramoyl-L-alanine amidase (41, 42), detected in one-week-old SBFs, is involved in this process, as weakening the link between the mycomembrane and peptidoglycan has been suggested to stimulate MV blebbing in the mycobacteria (38). Taken together, these findings may explain why more cytoplasmic proteins were detected on this biofilm subtype, as the maturing biofilm cells grow under reduced oxygen tension and anoxia is one of the factors able to trigger the membrane vesiculation.

Bacterial MVs are involved in i.e., cell-cell communication, biofilm formation, virulence, antibiotic resistance, iron scavenging, nutrient acquisition and modulating the host immune system (43). We detected several cytoplasmic proteins involved in signal transduction (e.g., PknL specific to SBFs and an adenylate cyclase detected only in biofilm-ECMs) and enzymes involved in biofilm formation. GroEL1 and Fatty-Acid-Synthase system (FAS-I and FAS-II) enzymes were among the detected proteins that coordinate biofilm formation in mycobacteria. The GroEL1 chaperone is involved in the synthesis of mycolic acids (MAs) that eventually become inserted in the mycomembrane as trehalose dimycolates (TDM) and monomycolates (TMM) beneath the capsule (14, 21). This chaperone interacts with ketoacyl-ACP synthase KasA (FAS-II) to modulate the synthesis of short-chain MAs specifically during biofilm formation (21). A lack of GroEL1 has been reported to prevent the biofilm formation and to affect the biosynthesis and composition of MAs in *Mycobacterium bovis* BCG, whereas the GroEL1 deficiency blocks the formation of mature biofilms in *M. smegmatis* (21, 24). In addition, the overexpression of KasA and the inactivation of other FAS-II enzymes, such as enoyl-ACP reductase (InhA) and 3-oxoacyl-[acyl-carrier-protein] synthase 2 (KasB), have also been reported to prevent biofilm formation and formation of cords by reducing the cyclopropanation of MAs (14, 21, 25). Here, GroEL1, KasA and InhA were detected as more abundant in the SBFs, implying that these enzymes could support the initial stages of SBF-type biofilm growth, as GroEL and KasA were detected with the highest abundancies already on the two-day-old SBFs.

Although no cell lysis was seen during the sample preparation for proteomic analysis **(Table S7)**, we cannot exclude the possibility that some of the cytoplasmic or inner-/myco-membrane-proteins were released by autolysis during growth. In other Gram-positive bacteria, cytoplasmic proteins reach the extracellular space via regulated autolysis (involving autolysins/peptidoglycan hydrolases), and, as soon as the pH of the culture medium drops (due to the active metabolism of the growing cells), many of the released proteins show an enhanced ability to bind to the cell wall and biofilm-ECM structures (43-48). SBF cells are exposed to hypoxic conditions, and oxygen limitation acidifies the biofilm matrix (48), allowing for a more efficient interaction between the cytoplasmic proteins and biofilm-ECM structures. Thus, this could explain the presence of r-proteins as the largest cytoplasmic protein group already on two-day-old SBFs; the strong positive charge of these proteins has been proposed to mediate electrostatic interactions with anionic cell-surface components, which promotes cell aggregation and biofilm stabilization (48). Since the exposed mycomembranes with MAs as the major components create a condition stimulating an interaction with many cytoplasmic proteins, pH-dependent binding with the cell surface components could also explain why cytoplasmic proteins were detected on Mmr cells grown on Tween-80.

### Biofilm subtypes differ in terms of secreted virulence and adhesion factors

The proteomics data indicated that the mycomembrane-associated PPE/PE family proteins were remarkably greater in number in the PL than in the PBFs or SBFs, indicating that Mmr in a single cell state could more readily interact with the host, and modulate the host immune response and/or nutrient transport (49, 50). PL cells were cultured in the presence of Tween-80, which, in detaching the mycobacterial capsule (17), most likely helped identify these immunogens. Tween-80 can also induce alterations in the morphology, pathogenicity, and virulence of mycobacteria (51). For example, genes encoding lipases and cutinases have been shown to be significantly upregulated in Mtb in response to this nonionic surfactant. Our data is in line with this by showing that several lipases/cutinases, with a likely ability to hydrolyze Tween-80, were more abundant on PL cells compared to biofilms. As Tween-80 is considered to mimic a lipid rich milieu of macrophages (51), the detected PL-proteome changes here may reflect a metabolic adaption to conditions faced in vivo.

Our findings also suggest that Mmr uses different T7SS pathways in SBFs and PBFs for virulence and adherence. For example, the Esx1-secretion components and substrates (EsxA/B, EspB, EspF, EccA1, EspG1, EspH, EspL and MycP) were detected as more abundant in the PBFs, while those associated with the Esx5-type secretion were overall more abundant in the SBFs (Ecc, EspG, PPE/PE proteins). Both secretion pathways can contribute to virulence and subverting the host immune system in Mtb (52). The major subtype-dependent differences between the PBFs and SBFs were related to invasion and adherence, including the MCE proteins, fibronectin binding APA and HphA, which can modulate host cell signaling as well as aid adhesion or entry into host cells (53-55). All these proteins were significantly more produced on the PBFs than the SBFs, and, in the case of MCEs, may also involve MVs, as these adhesins are located on the inner membrane of the mycobacterial cell wall. HphA also has implications in promoting cell-cell aggregation in Mtb (56), suggesting that this adhesin could also contribute to cording during PBF-type growth.

### Biofilm subtypes use different tolerance- and persistence-conferring mechanisms

Tolerance is defined as the extent of time that bacteria can survive in the presence of a high antibiotic concentration (31), whereas persisters are a subpopulation of phenotypically drug tolerant cells that do not grow in the presence of an antibiotic (32). We show that antibiotic killing of biofilm cells occurs at a significantly slower rate when compared to PL cells. The time-kill curve indicated the temporally increased formation of a persistent subpopulation with slower killing kinetics as well as the formation of persisters in SBFs already after one week. At this stage, Mmr biofilms remained metabolically active and replicating, indicating that persistence develops due to phenotypic differentiation during biofilm growth rather than via the induction of dormancy.

The proteomic findings suggest that Mmr could use both overlapping and subtype-specific mechanisms for increasing its tolerance and persistence, in which MVs or other non-classical routes for protein export may play a role. Here, most significant proteome changes related to cytoplasmic and inner-/mycomembrane-proteins and included enzymes/proteins involved in the TCA cycle and glyoxylate shunt, mycolic acid synthesis stress response, and energy and redox metabolisms. A recent transcriptome analysis of another non-tuberculous mycobacterial model, *Mycobacterium abscessus*, supports our findings; biofilm growth activated the glyoxylate shunt, redox metabolism and the MA synthesis-associated elongation and desaturation pathways. The TCA cycle associated enzyme CitA was recently reported to control the asymmetric cell division in *Caulobacter crescentus* (57). This process has been shown to lead to the formation of heterogenous cell populations in biofilms, macrophages, and granulomatous lesions also in mycobacteria (7, 58, 59). Here, our findings indicated the presence of this enzyme on one-week-old SBFs, suggesting that the asymmetric cell division occurs before the PBFs are formed. Moreover, arabinosyltransferases EmbA and EmbB, involved in the polymerization of the arabinogalactan, were also detected with high abundances in SBFs by one week onward, suggesting that strengthening the arabinogalactan could further help residing cells, including the persisters, increase their tolerance to rifampicin, as demonstrated with Mtb persisters under hypoxia (60). Taken together, these findings strengthen the hypothesis that persisters are indeed formed in one-week-old SBFs, and supports the results obtained with the biofilm killing assay on the SBFs at this time point.

We also suggest that cells in PBFs use different TCA cycle enzymes, such as aconitase (ACN), malate dehydrogenase (MDH), enolase (ENO), and/or fructose-bisphosphate aldolase (FBA), to maintain long-term survival. In other gram-positive bacteria these enzymes belong to known moonlighting proteins with established secondary roles outside of the bacterial cell (e.g., adhesion)(30). In mycobacteria, these enzymes have been reported to contribute to increased viability or persistence (61-63). The associated glyoxylate shunt could also be involved (64), as isocitrate lyase 1 (ICL1) was detected as more abundant on the SBFs, implying that this enzyme could help residing cells increase their antioxidant defense and antibiotic tolerance (65). In contrast, ICL2 was produced more on the PBFs, which may help the cells to survive under starvation conditions when fatty acids are used as the primary carbon source (66). This is in line with the temporally increased production of diacylglycerol *O*-acyltransferase (Tgs1) in PBFs, which can promote the accumulation of triacylglycerols (TAGs); a process that has been considered a hallmark feature of persisting Mtb/latent TB and a long-term energy source for Mtb and have been found in substantial amounts in the mycobacterial cell wall (67, 68). The detection of trehalase as significantly more abundant in four-to 12-week-old PBFs, strengthens the idea that cells within this biofilm subtype suffer from nutrient stress and activate trehalose salvage/recycling to promote redox and energy homeostasis, as seen under carbon limitations in Mtb (69). These findings may also explain the detection of proteases, chaperones, and assisting stress-proteins in high numbers in the biofilm-ECMs, including e.g., the proteases Clp/Lon and the cold-shock protein CpsD, with known implications in stringent response, persistence and/or post-antibiotic recovery (70-72). These proteins were detected here as more abundant in the PBFs than in the SBFs, implying that these pathways are preferred in PBFs to maintain viability.

A recent study comparing high numbers of persister Mtb mutants using genomics and transcriptomics indicated a significant upregulation of energy production pathways and pathways involved in redox reactions (oxidoreductase) (73). The ECM-proteome changes occurring during SBF-type growth are in line with this report, as the components of the respiratory electron transfer chain (cytochrome bc1 complex, cytochrome c terminal oxidase, and F0F1-ATPase synthase) were detected as more abundant on the SBFs facing more hypoxic conditions than the PBFs. Our findings also agree with previous reports showing that the electron transfer chain is essential for maintaining ATP homeostasis and the viability of nonreplicating/persistent Mtb cells under hypoxia (74-76). In addition, we show that both redox and iron metabolism could also play a biofilm subtype-specific role in helping the cells cope with hypoxia/aeration-related stress (77); several oxidoreductases, thioredoxin and a superoxide dismutase (SOD) were overall more abundant in the SBFs, and a catalase-peroxidase (KatG) and alkyl hydroxyperoxidases (AhpCF) as more abundant in the PBFs. These enzymes have been shown to protect Mtb against oxidative stress by the reduction of superoxide radicals into less toxic intermediates for inhibiting autophagy, apoptosis and cellular damage (78). Iron storing proteins ferritin (BfrB) and bacterioferritin (BfrA) can confer increased redox resistance on Mtb and protect the cells against oxidative stress and hypoxia, respectively (79). Here, these iron storing proteins displayed biofilm subtype-specific abundance changes, implying that SBFs could rely on BfrB to cope with hypoxia and PBFs on BrfA to help cells grow at the air–liquid interface.

## CONCLUSIONS

The present study reports an in-depth view of ECM-proteome changes occurring in Mmr ATCC927 during biofilm growth in vitro from two days to three months. We show that this non-tuberculous mycobacterial model forms SBFs already after two days, whereas the formation of detectable PBFs was observed after two weeks of growth in the absence of Tween-80. Both biofilm subtypes were formed physically under the same conditions with clearly distinct growth morphologies: SBFs with lichen-like structures and PBFs with ribbon-like cords. We show that the reduced CelA1-mediated cellulose hydrolysis is necessary to establish proper biofilm growth, growth morphology and increased tolerance to rifampicin for both biofilm subtypes. The formation of persisters in both biofilm subtypes and increased tolerance was further confirmed by the newly established bioluminescence-based time-kill assay, which provides an effective tool for quantifying tolerance and persistence in Mmr. The proteomic findings imply that subtype-dependent changes in the MA synthesis and modification, Esx1-type secretion, and the production of specific adhesins were the major drivers of distinct biofilm growth morphologies. We also propose that pathways associated with MA biosynthesis, development of tolerance/persistence and oxidative/redox stress are differentially used in PBFs and SBFs to maintain prolonged viability. Possible explanations for these differences include the different oxygen tensions encountered by the biofilm subtypes, differences in membrane vesiculation activities and/or other non-classical pathways for protein export. Taken together, this is the first study reporting on ECM-proteome dynamics in maturing mycobacterial biofilms and predicting biofilm subtype-specific changes in cell-cell communication, biofilm matrix formation, virulence, and tolerance/persistence. This is also the first time that the kinetics of persistence have been explicitly measured from mycobacterial biofilms.

## MATERIALS AND METHODS

### Preparing bacterial cells for surface proteomics

*Mycobacterium marinum* (ATCC 927) with the pTEC27 plasmid expressing the red fluorescent protein tdTomato (Addgene #30182, http://n2t.net/addgene:30182) (29) was pre-cultured on Middlebrook 7H10 plates with 10% (v/v) Oleic Albumin Dextrose Catalase (OADC) enrichment (Fisher Scientific, NH, USA) and 0.5% (v/v) glycerol at 29 °C for one week. For planktonic cultures, an inoculum of Mmr was transferred into a Middlebrook 7H9 medium supplemented with 10% (v/v) ADC (Fisher Scientific, NH, USA), 0.2% (v/v) glycerol, and 0.2% (v/v) Tween-80 (Sigma-Aldrich, MO, USA), and the cells were cultured at 29 °C in cell culture flasks with filter caps. After three days of incubation the cell cultures were diluted to obtain an OD_600_ of 0.042 and the dilutions were cultured for an additional 2 days at +29 °C until harvesting. For the biofilm cultures, a Middlebrook 7H9 medium with the ADC growth supplement but without Tween-80 or glycerol was used. The inoculum was cultured for three days at +29 °C until the OD_600_ reached 0.45. The cell cultures were diluted 1:40, and the dilutions were divided into 10 ml aliquots. The cap of each tube was sealed with Parafilm M® laboratory wrapping film, and the cultures were incubated at +29 °C. Planktonic and biofilm cell samples (SBFs and PBFs separately) were collected at the time points indicated in **Figure 2A**. All the cultures were performed in quadruplicates. Planktonic cells were harvested by centrifugation (3 min, 5 000g, +4 °C) and the pelleted cells were suspended gently in ice-cold buffer (100 mM Bis-Tris, pH 6.5) to remove interfering/non-specifically bound proteins. This step prevents the detachment/removal of cytoplasmic moonlighters bound to the cell surfaces/biofilm-ECM (43-46, 79, 80). The PBFs were collected with an inoculation loop, the extra medium was removed by pipetting to avoid cross-biofilm type contamination, and the SBFs were harvested by pipetting/scraping. The PBFs and SBFs were collected in separate Eppendorf tubes the ice-cold buffer (100 mM Bis-Tris, pH 6.5). Cells (planktonic and biofilm cultures) were pelleted by centrifugation (3 min, 5 000*g*, +4 °C) and the washed cells were suspended gently in 95 μL of 100 mM TEAB (triethylammonium bicarbonate, pH 8.5) for the enzymatic shaving reaction.

### Trypsin/Lys-C shaving of planktonic and biofilm cells

Peptides from cell-surface/biofilm-ECM-associated proteins were released via a Trypsin/Lys-C mix (Promega) at a final concentration of 50 ng µL^-1^, and the digestions were incubated at 37 °C for 20 minutes. The method was validated by counting the number of colonies formed on the planktonic/single and biofilm cells treated with/without the enzyme mix **(Table S7)**. The released peptides and the enzymes were recovered by filtration through a 0.2 μm acetate membrane (Costar® Spin-X Centrifuge Tube Filter, Corning Inc., Corning, NY, US) by centrifugation (8000*g*, 3 min, 20 °C). Flow-troughs were incubated for 16 hours at 37 °C. The concentration of released peptides in each sample was measured with a NanoDrop2000 spectrophotometer (Thermo Scientific). Digestions were terminated with 0.6% (v/v) trifluoroacetic acid (TFA) (Sigma Aldrich) and the peptides were purified using ZipTip C18 (Millipore) according to the manufacturer’s instructions and dried using a miVac centrifugal vacuum concentrator (GeneVac).

### LC-MS/MS analysis

The peptides were dissolved in 0.1% (v/v) formic acid (FA) and analyzed with nanoLC-MS/MS using an Easy-nLC 1000 nano-LC system (Thermo Scientific) coupled with a quadrupole Orbitrap mass spectrometer (Q Exactive™, ThermoElectron, Bremen, Germany) as previously reported (80). The obtained MS raw data was processed via MaxQuant software (version v.1.6.1.0) with the built-in search engine, Andromeda (81, 82), using a protein database comprising all 5,564 Mmr protein sequences (Uniprot proteome: up000257451, genome accession: PEDF01000000) both forward and reverse. Carbamidomethyl (C) was set as a fixed and methionine oxidation was set as a variable modification. Tolerance was set to 20 ppm in the first search and 4.5 ppm in the main search. Trypsin without the proline restriction enzyme option and with two allowed miscleavages was used. The minimal unique plus+ razor peptide number was set to 1; the FDR was set to 0.01 (1%) for peptide and protein identification; and LFQ with default settings was used. The mass spectrometry proteomics was deposited in the ProteomeXchange Consortium via the PRIDE (83) partner repository with the dataset identifier PXD02010.

### Proteome statistics and bioinformatics

The identified Mmr proteins were manually curated by characterizing hypothetical and tentatively annotated proteins with the aid of the Basic Local Alignment Search Tool (BLAST) program from the National Center for Biotechnology Information (NCBI) (84-86), combined with CDD/SPARCLE conserved domain identification (87) and SmartBLAST (UniProt) searches. General protein functions were annotated using the Gene Ontology (GO) database (88). Isoelectric points (pIs) and molecular weights (MWs) for the identified proteins were predicted using EMBOSS Pepstats (89) at https://www.ebi.ac. uk/Tools/seqstats/emboss_pepstats/. The presence of possible protein secretion motifs (classical and nonclassical) for all the predicted and identified proteins was obtained with SignalP4.1 (90) (http://www.cbs.dtu.dk/services/SignalP/) and SecretomeP 2.0/SecP (91) (http://www.cbs.dtu.dk/services/SecretomeP/). The presence of transmembrane spanning domains/helices (TMDs) was determined with the TMHMM Server v. 2.0 at http://www.cbs.dtu.dk/services/TMHMM/ (92, 93) for the identified proteins.

For indicating statistically significant abundance changes the log2-transformed LFQ data was analyzed in Perseus v.1.6.2.3 (94) using a Student’s *t*-test with permutation-based FDR adjustment. For the multivariate analyses, the missing values were replaced by imputed values from the normal distribution (width = 0.3, down shift = 1.8) and then normalized (z-score) prior to ANOVA for multi-sample testing (S0 set to 0.1 and a permutation-based FDR of 5%) and hierarchical clustering/PCA. STRING Protein Interaction Network and Functional Enrichment Analyses (GO, KEGG, InterPro, Pham) were studied using the STRING database v. 11 (95). Interaction scores were set to high (0.700) confidence, and the interacting proteins were clustered using Markov clustering (MCL) with the inflation parameter set to 4.0–6.0. Functional enrichments were statistically assessed with both rank- and gene set-based approaches (FDR of 0.05).

### Creation of the CelA1 overexpression construct in Mmr

The Mmr CelA1 overexpression strain was created by ordering the MMAR_0107 open reading frame in the pUC57 vector with appropriate restriction sites from GenScript and subcloning the construct into the pTEC27 vector (AddGene) (29), which carryies the red fluorescent protein tdTomato. The sequence of the plasmid was confirmed by sequencing. The resulting plasmid was transformed into an electrocompetent Mmr ATCC927 strain. Transformants were selected on Middlebrook 7H10 agar plates containing 10% (v/v) OADC enrichment, 0.5% (v/v) glycerol and 75 µg mL^-1^ hygromycin.

### RNA and DNA extractions

For RNA and DNA extractions, the *CelA1* overexpression strain and Mmr were precultured on MiddleBrook 7H10 plates and transferred into the Middlebrook 7H9 medium described above (75 µg mL^-1^ hygromycin for the CelA1 strain). After three days, the bacterial cells were harvested, pelleted, and homogenized in TRI Reagent (Thermo Fisher Scientific, NH, USA) with ceramic beads using the PowerLyzer24 (Mobio, CA, USA). After homogenization, the samples were sonicated for nine minutes and the RNA and DNA were extracted according to the manufacturer’s instructions.

### *CelA1* expression and the quantification of mycobacterial loads by qPCR

Prior to qPCR analysis, RNA was reverse transcribed into cDNA with a Reverse Transcription kit (Fluidigm, CA, USA) according to the manufacturer’s instructions. *CelA1* expression was measured using soFast EvaGreen Supermix with the Low ROX qPCR kit (Bio-Rad, CA, USA) and the CFX96 qPCR system (Bio-Rad, CA, USA). The primers used for *CelA1* were: (forward: 5’-ACACTCCGCAGTCCTACT-3’ and reverse: 5’-TAGAGCGTC AGAATCGGC-3’). The number of mycobacterial cells in the sample was quantified using the SensiFAST SYBR No-ROX qPCR kit (Bioline, London, UK) on bacterial DNA according to the manufacturer’s instructions. The primers used for Mmr quantification were targeted against 16S-23S, locus AB548718 (forward: 5’-CACCACGAGAAACACTCCAA-3’ and reverse: 5’-CACCACGAGAAACACTCCAA-3’). Each bacterial quantification qPCR run included a standard curve of the known amounts of Mmr DNA. The mycobacterial cell number in each sample was used to normalize the *CelA1* expression.

### Widefield deconvolution microscopy (WDeM) of Mmr biofilms

PBFs and SBFs formed by Mmr with pTEC27 (WT), expressing the red fluorescent protein tdTomato (29), and Mmr overexpressing CelA1 were prepared as follows. Briefly, the cells were incubated at 29 °C and the surface-attached cells were imaged at seven, 14 and 21 days after dilution. In situ imaging of the SBFs was conducted with Nikon FN1 upright epifluorescence microscope equipped with 20x/0.8 dry objective, Hamamatsu ORCA-Flash4.0 V3 Digital CMOS camera and CoolLED pE-4000 light source. tdTomato was excited with 550 nm LED and fluorescence was collected with 617/73 band-pass emission filter. Image stack were collected with 2µm intervals (x-y pixel size 325 nm). The data was deconvolved with Huygens Essential deconvolution software (SVI, Amsterdam, Netherlands) using 200 iteration limit, signal-to-noise ratio of 30 and quality threshold of 0.01.

### Biofilm tolerance assays

The role of CelA1 overexpression in the antibiotic tolerance of Mmr was assessed as follows. First, the *CelA1* overexpression strain and pTEC27 control strain were cultured for one week on 7H10 plates (10% OADC, 0.5% glycerol + 75 µg mL^-1^ hygromycin) and then transferred in a Middlebrook 7H9 medium supplemented with 10% ADC and 75 µg mL^-1^ hygromycin) at an OD_600_ of 0.1 to initiate biofilm growth. Aliquots of bacterial suspension (192 µl of per well) were added to 96-well-plates in triplicate, sealed with parafilm and incubated at +29 °C in the dark. Planktonic cultures grown in the presence of 0.2% glycerol were used as controls. Eight µl of antibiotics per well was added two, four, seven and 14 days after the start of the liquid culture. The final antibiotic concentrations used were 400, 100, 25 and 6.25 µg mL^-1^ for the Rifampicin TOKU-E solution. Untreated wells were used as controls. Ten µl per sample was plated on 7H9 plates (10% OADC, 75 µg mL^-1^ hygromycin) one week after the addition of antibiotics. The plates were incubated at +29 °C for seven to nine days and the colonies were counted.

### Biofilm persistence assays

Mmr (ATCC 927) with a bioluminescence cassette in the pMV306hsp+LuxG13 plasmid was used for antibiotic tolerance assays. pMV306hsp+LuxG13 was provided by Brian Robertson and Siouxsie Wiles (Addgene plasmid #26161; http://n2t.net/addgene:26161). To measure the kinetics of bacterial killing, the bioluminescent Mmr strain was first cultured on Middlebrook 7H10 agar (Sigma-Aldrich) supplemented with 0.5% (v/v) glycerol (Sigma-Aldrich) and 10% (v/v) OADC enrichment (BD, Becton Dickinson) at 29 °C for seven days in the dark. To initiate biofilm formation, the Mmr cells were suspended in Middlebrook 7H9 broth (Sigma-Aldrich) supplemented with 10% (v/v) ADC enrichment (BD, Becton Dickinson) at a starting OD_600_ of 0.1. Planktonic cultures were prepared in the same way except that the medium contained 0.2% (v/v) glycerol (Sigma-Aldrich) and 0.2% (v/v)Tween-80 (Sigma-Aldrich). Bacterial suspensions (192 µL per well in triplicates) were divided on to white 96-well plates (Perkin Elmer). The biofilm cultures were sealed with laboratory film and incubated at 29 °C in the dark to the desired ages. Rifampicin solution (TOKU-E) in water at a final concentration of 400 µg mL^-1^ corresponding to 64 x MIC (minimum inhibitory concentration) was added to the bacterial suspensions and incubated for seven days at 29 °C in the dark. The bioluminescence signal was measured with an EnVision plate reader (Perkin Elmer) as a readout for bacterial survival three times for three seconds per well daily from a white 96-well plate for seven days. The background signal from media only wells was first subtracted from the sample wells and an average of the three measurements normalized with the starting bioluminescence signal was used to draw time-kill curves of the bacterial population in the biofilms at different maturation stages.

To compare the level of persistence/tolerance in the PBFs and SBFs, Mmr was cultured in a total volume of 10ml at the starting OD_600_ value of 0.1. After two weeks, the biofilms were collected separately from the tubes by lifting the pellicle with a 1-µL inoculation loop coupled with careful pipetting. The pellicle and submerged biofilms were centrifuged at 10,000*g* for three minutes, the supernatants were collected, and the wet weight of the bacterial mass was measured. The bacterial cells were suspended into previously collected spent media at the concentration of 15 mg mL^-1^, vortexed briefly, and divided on white 96-well plates (Perkin Elmer) with 192 µL of cell suspension per well in triplicate. Eight µl of TOKU-E solution at a final concentration of 400 µg mL^-1^ were pipetted on the bacterial suspension. Liquid cultures were incubated for seven days at 29 °C in the dark and the bioluminescence signal was measured daily with an EnVision plate reader (Perkin Elmer) three times for three seconds per well. The background signal from the media-only wells was first subtracted from the sample wells, and an average of the three measurements was normalized with the starting bioluminescence signal measured just before adding the rifampicin. The statistical significance of the differences between the time-kill curves was tested with a log-rank test using Prism5 software (GraphPad).

## ACKNOWLEDGEMENTS

The financial support of the Academy of Finland (projects no. 322010 (AF, JYK, MP) and 326674 (MP), Jane and Aatos Erkko Foundation (MP), Sigrid Jusélius Foundation (MP), Foundation of the Finnish Anti-Tuberculosis Association (Suomen Tuberkuloosin Vastustamisyhdistyksen Säätiö)(HL, LMV, KS) and Tampere Tuberculosis Foundation (MP, KS, HM, HL, LMV) is greatly appreciated. Mass spectrometry-based proteomic analyses were performed by the Proteomics Core Facility, Department of Immunology, University of Oslo/Oslo University Hospital, which is supported by the Core Facilities program of the South-Eastern Norway Regional Health Authority. This core facility is also a member of the National Network of Advanced Proteomics Infrastructure (NAPI), which is funded by the Research Council of Norway INFRASTRUKTUR-program (project number: 295910)(TAN).

## SUPPLEMENTAL MATERIAL

**Figure S1. (A)** Pellicle-type (PBF) and submerged (SBF) biofilms after culturing for two weeks (left) and 12 weeks (right). **(B)** Distribution of overlapping protein identifications within four replica samples from planktonic cell surfaces and biofilm-ECMs. PL_log, logarithmic planktonic cells; SBF, submerged type biofilms; PBF, pellicle type biofilms.

**Figure S2. (A)** Distribution of identified proteins in terms of their predicted secretion motifs and the number of predicted TMDs. PL_log, logarithmic planktonic cells; PBF_all and SBF_all, all identified proteins from pellicle and submerged biofilm matrices, respectively. Other, proteins without any known motifs for classical or non-classical secretion. **(B)** Venn diagrams indicating the core and marker proteomes within all identifications (planktonic and biofilms) at each time point.

**Figure S3. (A)** The bioluminescence-based readout of the biofilm killing assay was validated using the OD_600_–method to monitor bacterial growth at varying time points of growth, showing similar killing kinetics as observed with the bioluminescence measurements of the same samples. Here, the four-day-old biofilm and two-day-old planktonic cells were treated with 200 μg mL^-1^ rifampicin. The mean of three biological replicates of the OD_600_-based assay is shown. **(B)** Mmr growth is accompanied by increased bioluminescence values in maturing biofilms without the antibiotic treatment, even at timepoints of over one week of biofilm culture. The bioluminescence was measured three times/three seconds using EnVision equipment (Perkin Elmer), and the mean of relative light units (RLUs) per one second was calculated. Means and SEMs of three biological replicates are shown.

**Table S1**. List of all proteins identified on Mmr cells during different growth modes. The colored cells refer to the average (log2) raw intensity values for proteins detected in at least two replica samples. Cells in grey, protein was not detected. PL_log, logarithmic planktonic cells; PBF, pellicle type biofilm cells; SBF, submerged type biofilm cells.

**Table S2**. Proteins specific to pellicle biofilms (PBFs), submerged biofilms (SBFs) and to both biofilms that lack an identifiable counterpart on planktonic cell surfaces. Proteins detected in each replica samples are shown. Color gradient bar refers to log2-transformed raw intensity values (avr., n ≥ 2): blue, low abundance; yellow, high abundance; PBF, pellicle biofilm; SBF, submerged biofilm.

**Table S3**. Log2-transformed MaxLFQ data with minimum of two valid identifications (out of four) in at least one group, and statistically significant protein abundance changes between the PL and PBF_2, PL and SBF-2d, PBF_2w_SBF_2d, PBF_12w and SBF_12w, PBF_4w and 12w and SBF_4w and 12w.

**Table S4**. Statistically significant protein abundance changes within planktonic cell surfaces and biofilm-ECMs. Colored cells in column “Cluster” correspond to those used in the heat-map (Fig. 5). Significant changes were calculated using a multiple-sample test (ANOVA model, FDR < 0.05, S0 = 0.1). Color intensity code bar below, blue - low abundance; yellow - high abundance.

**Table S5**. Functional enrichment analysis (GO, KEGG, InterPro, Pham) on cluster 1, 2 and 6 proteins (Figure 5) were studied using the STRING database v. 11 with both the rank- and gene set-based approaches (FDR of 0.05).

**Table S6**. Key proteome changes within the planktonic cell surfaces and biofilm matrices at different time points of growth. Gradient bar, normalized identification intensity values (avg. ≥ 3).

**Table S7**. Colony ability of non-shaved and enzymatically shaved biofilm and planktonic cells. Biofilm cells were cultured for two weeks and planktonic cells for two days, as described in material and methods. Cells in three biological replicates were suspended in trypsin/Lys-C digestion buffer and then divided into two aliquots; the first aliquot was taken as the non-shaved cell control containing only the digestion buffer, and the second aliquot of cells was treated with the trypsin/Lys-C enzyme. After 20 min incubation at 37 °C, the cell suspensions were suspended gently in PBS containing 0.2% Tween80 (v/v) and serially diluted cells were spotted in four technical replicates (10 uL each) on an agar plate. After one week cultivation at 37 °C, the colonies were calculated and compared under different conditions. nd, colonies could not be counted due to the presence of cell aggregates.

